# Direct gamete sequencing reveals no evidence for segregation distortion in house mouse hybrids

**DOI:** 10.1101/008672

**Authors:** Russell Corbett-Detig, Emily Jacobs-Palmer, Daniel Hartl, Hopi Hoekstra

## Abstract

Understanding the molecular basis of species formation is an important goal in evolutionary genetics, and Dobzhansky-Muller incompatibilities are thought to be a common source of postzygotic reproductive isolation between closely related lineages. However, the evolutionary forces that lead to the accumulation of such incompatibilities between diverging taxa are poorly understood. Segregation distorters are believed to be an important source of Dobzhansky-Muller incompatibilities between *Drosophila* species and crop plants, but it remains unclear if these selfish genetic elements contribute to reproductive isolation in other species. Here, we collected viable sperm from first-generation hybrid male progeny of *Mus musculus castaneus* and *M. m. domesticus*, two subspecies of rodent in the earliest stages of speciation. We then genotyped millions of single nucleotide polymorphisms in these gamete pools and tested for a skew in the frequency of parental alleles across the genome. We show that segregation distorters are not measurable contributors to observed infertility in these hybrid males, despite sufficient statistical power to detect even weak segregation distortion with our novel method. Thus, reduced hybrid male fertility in crosses between these nascent species is attributable to other evolutionary forces.

## Introduction

The Dobzhansky-Muller model [1,2] is widely accepted among evolutionary biologists as a primary explanation for the accumulation of intrinsic reproductive incompatibilities between diverging lineages [3,4]. Briefly, this model posits that genes operating normally in their native genetic background can be dysfunctional in a hybrid background due to epistatic interactions with alleles from a divergent lineage. Although elucidating the molecular basis of speciation has been a central focus for decades, loci contributing to Dobzhansky-Muller incompatibilities (DMIs) have proved challenging to identify primarily because they are, by definition, incompatible in combination (review by [3–6]). As a result, the specific genetic changes responsible for the onset of reproductive isolation between lineages remain largely obscure.

The rapid evolution of selfish genetic elements within lineages is thought to be a potent source of DMIs between diverging taxa. Segregation distorters are one such selfish element that increase their transmission through heterozygotes by either disabling or destroying gametes that failed to inherit the distorting allele [7,8]. Because males heterozygous for a distorter produce fewer viable sperm, segregation distorters can decrease the fitness of carriers. In this case, other loci in the genome are expected to evolve to suppress distortion [9]. This coevolution of drivers and suppressors has been suggested to be a widespread source of DMIs between diverging lineages: hybrids of isolated populations in which such coevolutionary cycles have occurred may suffer lower fertility as drivers become uncoupled from their suppressors in a mixed genome [10–12]. Indeed, there is evidence that SDs contribute to hybrid male sterility in several *Drosophila* species pairs (*e.g.* [13–15], reviewed by [4,12]) as well as in many crop species (*e.g.* [16–20]). However, comparatively little is known about genetics of speciation aside from these groups, and it remains unclear if distorters contribute to hybrid sterility in other taxa more generally.

Analyses aimed at identifying the genetic targets of positive selection suggest that segregation distorters may be an important source of DMIs in mammalian lineages. One particularly intriguing finding shows a substantial overrepresentation of loci associated with spermatogenesis and apoptosis within the set of genes with the strongest evidence for recurrent positive selection in mammals (*e.g.* [21,22]). These functions in turn are potentially driven, at least in part, by segregation distorters, which are expected to leave just such a mark of selection as they sweep through a population. Therefore, mammals are an appealing group in which to test for segregation distortion and its role in speciation.

In particular, *Mus musculus domesticus* and *M. m. castaneus* are two subspecies of house mice in the earliest stages of evolving reproductive isolation [23,24]. Indeed, these subspecies are estimated to be approximately 500,000 years diverged from one another [24]. Hybrid males suffer from reproductive deficiencies [25]; specifically, the vas deferens of first-generation hybrid (F_1_) males contain more apoptotic sperm cells than either pure strain, and numerous loci affecting fertility in hybrid males have been reported, particularly in F_2_ individuals [26]. Finally, Wager [27] identified eight genomic regions that exhibited significant deviations from Mendelian segregation in an F_2_ mapping population derived from these two subspecies, which may be consistent with the action of segregation distorters in their hybrids (but see below). In combination with the comparative genomic evidence and phenotypic observations described above, these data suggest that coevolution of SDs and their suppressors may contribute to DMIs in *M. musculus.*

The conventional approach to identifying SD relies on detecting a skew in the allele frequencies of second-generation hybrids in a large genetic cross. However, methods that rely on genotyping progeny unavoidably conflate segregation distortion, female effects on sperm function, and differential viability. Additionally, practical issues limit the power of these experiments—specifically, the ability to produce and genotype hundreds to thousands of individuals in order to detect distorters of small effect—particularly in vertebrates. Therefore, as a result of modest sample sizes, many experiments designed to detect SD using genetic crosses are underpowered and unable to detect even moderate distortion.

Here, we explore a novel approach to surveying the genome for SD by directly sequencing viable gametes from F_1_ hybrid *M. m. domesticus/M. m. castaneus* males. Briefly, we enriched for viable sperm in hybrids and then sequenced these sperm in bulk, along with a control tissue, to identify any skew in the representation of either parental chromosome in the viable sperm relative to the control (Figure 1). While we demonstrate via simulation that our experimental design has excellent power, we find no evidence of SD in this cross, suggesting that segregation distorters are not a primary contributor to male infertility in *M. m. castaneus* an d*M. m. domesticus* hybrids. Nonetheless, this approach can be applied to a wide range of species, and we therefore expect that it will be a useful means to study the frequency and impact of segregation distortion more generally.

**Figure 1.**
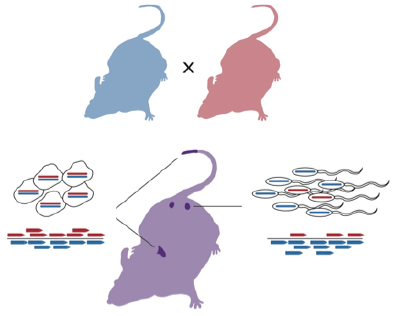
Schematic of experimental cross scheme. Inbred parental strains were crossed, and individual F1 males (purple) sacrificed at between 3 and 6 months, when their sperm were subjected to a swim up assay. Libraries were prepared from liver or tail (control; left) and sperm (experiment; right) samples, sequenced, and then aligned to a diploid reference genome; subspecies of origin were determined for as many sequences as possible.

## Materials and Methods

### Reference Genome Assembly

To generate robust genome assemblies for each of the two strains of interest, we aligned all short read data for *M. m. castaneus* strain (CAST/EiJ) and *M. m. domesticus* strain (WSB/EiJ) from a recent large-scale resequencing project [28] to the MM9 genome assembly using BWA v0.7.1 [29] for initial mapping. For reads that failed to map with high confidence, we remapped using stampy v1.0.17 [30]. We realigned reads that overlap indels, and called SNPs and indels for each strain using the Genome Analysis Tool Kit (GATK, [31]). For each program, we used default parameters, except that during variant calling we used the option ‘–-sample_ploidy 1,’ because the strains are extremely inbred.

We generated a consensus sequence for each strain at sites where both assemblies have high quality data. That is, if both CAST and WSB assemblies had a q30 minimum quality genotype (either indels or SNPs) that site was added to both consensus sequences. Otherwise, if either or both assemblies were below this quality threshold at a given site, we used the MM9 reference allele for both.

### Alignment Simulation

Our goal was to align short read data to a single diploid reference genome, comprised of assemblies from the two parental strains. The mapping quality, which indicates the probability that a read is incorrectly mapped in the position indicated by the aligner, should then provide a reliable means of distinguishing whether a read can be confidently assigned to one of the parental genomes. To confirm the accuracy of this approach and to identify suitable quality thresholds, we performed simulations using SimSeq (https://github.com/jstjohn/SimSeq). We used the sequencing error profiles derived from our mapped data (below) and found qualitatively similar error rates using the default error profile included with the SimSeq software package (data not shown). For both the CAST and WSB genomes, we simulated 10,000,000 pairs of 94-bp paired-end reads, whose size distribution was set to match that of our libraries (below). We then mapped these reads back to the single reference genome containing both CAST and WSB consensus sequences. We scored reads as ‘mapping correctly’ if they mapped to within 10 bp of their expected location measured by their left-most coordinate and on the correct subspecies’ chromosome. If the pair mapped, we required that the insert length be less than 500 bp, which is well within three standard deviations of the mean insert size of our data and should therefore encompass the vast majority of read pairs. If both reads in a pair mapped and met our criteria above, we used the higher mapping quality of the two, and discarded the other read. This filter is important, here and below, as it avoids counting pairs as though their provenance is independent of their pair.

### Experimental Crosses and Swim-Up Assay

To create first-generation (F_1_) hybrids of *Mus* subspecies, we crossed 2 *M. m. castaneus* males to 3 *M. m. domesticus* females and 2 *M. m. domesticus* males to 5 *M. m. castaneus* females in a harem-mating scheme. In total, we produced 8 male F_1_s in each direction of the cross. F_1_ males whose sire was *M. m. castaneus* (CAST genome) are referred to as CW, and those whose sire was *M. m. domesticus* (WSB genome) as WC. All males were housed individually for a minimum of two weeks prior to sacrifice between 90 and 120 days of age.

To enrich for viable sperm from each F_1_ male, we performed a standard swim up assay [32]. First, immediately following sacrifice, we collected and flash-froze liver and tail control tissues (liver samples, *N* = 16; tail samples *N* = 8). Then, we removed and lacerated the epididymides of each male, placed this tissue in 1.5 ml of human tubal fluid (Embryomax^®^ HTF, Millipore), and maintained the sample at a constant 37 °C for 10 minutes. Next, we isolated the supernatant, containing sperm that swam out of the epididymides, and spun this sample for 10 minutes at 250 g. We then discarded the supernatant, repeated the wash, and this time allowed sperm to swim up into the solution for an hour to select the most robust cells. Finally, we removed the solution, transferred them to new vial, pelleted these sperm by centrifugation, and froze them at -80 °C.

### Library Preparation and Sequencing

For each F_1_ hybrid male, we first extracted DNA from sperm, liver, and tail tissues identically using a protocol designed to overcome the difficulty of lysing the tightly packed DNA within sperm nuclei (Qiagen *Purification of total DNA from animal sperm using the DNeasy Blood & Tissue Kit; protocol 2*). We sheared this DNA by sonication to a target insert size of 300 bp using a Covaris S220, then performed blunt-end repair, adenylation, and adapter ligation following the manufacturer protocol (New England BioLabs). Following ligation, libraries were pooled into two groups of 16 and one group of 8 based on the adapter barcodes. Prior to PCR, each pool was subject to automated size selection for 450-500 bp to account for the addition of 175 bp adapter sequences, using a Pippen Prep (Sage Science) on a 2.0% agarose gel cassette. PCR was performed using six amplification cycles, and then we re-ran the size selection protocol to eliminate adapter dimer prior to sequencing. Finally, we pooled the three libraries and sequenced them on two lanes of a HiSeq 2500. Each sequencing run consisted of 100 bp paired-end reads, of which the first 6 bp are the adapter barcode sequence, and the remaining 94 bp are derived from randomly-sheared gDNA.

### Alignment and Read Counting

We aligned read data to the combined reference genome using ‘BWA mem’ as described above in the alignment simulation. We removed potential PCR duplicates using Picard v1.73. We then filtered reads based on the alignment filtering criteria described above for the simulated data. Because copy number variations may pose problems for our analysis, we attempted to identify and exclude these regions. Specifically, we broke the genome into non-overlapping 10 kb windows. Then, within each library, we searched for 10 kb regions that had a sequencing depth greater than two standard deviations above the mean for that library. All aberrantly high-depth windows identified were excluded in downstream analyses in all libraries. These regions, representing approximately 7% of the windows in the genome, are reported in Supplemental Table S1. Although deletions in one parental strain relative to the MM9 genome could also skew the parental allele frequencies for sequenced tissues, these copy number variable regions would affect both somatic tissues and gametes equivalently, and we therefore do not expect copy number variable regions to yield false positive results.

Next, to identify regions showing evidence of segregation distortion, we conducted windowed analyses with 1 Mb between the centers of adjacent windows. We counted reads in each window as a decreasing function of their distance from the center of the window, and included no reads at distances greater than 20 cM, thereby placing the most weight in a window on the center of the window. We then analyzed each window in two mixed-effects generalized linear models. Both models included random effects for the libraries and individuals. The first model includes no additional factors. The second had fixed effects for tissue, direction of cross, and an interaction term based on tissue by direction of cross effects, and thus has five fewer degrees of freedom than the first model. Hence, for each window, we assessed the fit of the second model relative to the first using a likelihood ratio test, wherein the log likelihood ratio should be chi-square distributed with 5 degrees of freedom. Afterwards, we applied a false-discovery rate multiple testing correction to the data [33]. We performed all statistical analyses in R [34].

### Power Simulations

To estimate the power of our method, we simulated distortion data. We began by selecting sites randomly distributed across the genome, and for each site drew a distortion coefficient from a uniform distribution between -0.05 and 0.05. Each read on the parental genome that was susceptible to distortion was counted on the distorting genome with probability equal to the distortion coefficient multiplied by the probability that no recombination events occurred between the distorted locus and the read. We also did the alternative (*i.e.* switching reads from the distorted against genome to the distorting genome) by multiplying by the probably that a recombination event was expected to occur in the genomic interval between the distorter and the read. We determined recombination probabilities using the genetic map reported in [35]. We performed the simulation for both parental genomes, and then again for each parental genome but with the distortion limited to one direction of the cross (*e.g.* only sperm from CW males experienced distortion). A direction-specific effect could occur if, for example, suppressing alleles are present on the Y chromosome of one subspecies and therefore are only present in CW or WC males.

## Results

After addressing the possibility of contamination, labeling, and quality issues (see Supplemental Text S1, Supplemental Table S2), we ran our analysis of the data across all autosomes, excluding regions with evidence for copy-number variations (described in Methods). With the exception of windows on chromosome 16 (see below), we found no windows with a statistically significant signature of segregation distortion. The lowest uncorrected *p*-value for any window (aside from those on chromosome 16) was 0.0224, which is not significant when we corrected for multiple tests. Thus, we did not find evidence for segregation distortion in any of the genomic regions considered (Figure 2).

**Figure 2.**
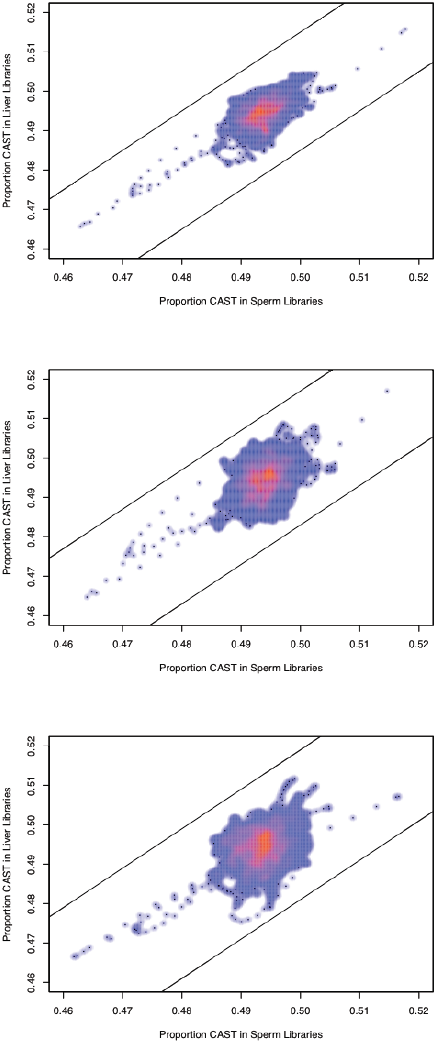
Average proportion CAST reads in sperm libraries versus liver libraries, using all males (A), using only CW males (B), and using only WC males (C). Lines indicate the approximate threshold at which we would have 50% power to detect distortion at the alpha = 0.0001 level (see Methods for how this threshold was calculated).

By contrast, on chromosome 16, we identified 15 contiguous windows with significantly skewed allele frequencies following correction for multiple comparisons (minimum *p* = 5.026E-4; Figure 3). However, upon closer examination, it appears that this signal is driven almost entirely by a single liver sample, that of individual CW10. If this sample is removed from the dataset, this chromosome no longer shows significant deviation from expectations. When comparing the relative read depths across chromosomes 16 and 1, CW10’s liver sample also appears to have disproportionately lower depth on this chromosome relative to CW10’s sperm sample (*p* = 3.02E-5; X^2^-test). These results suggest that this pattern is likely driven by a somatic aneuploidy event in CW10’s liver that occurred relatively early in liver development and is not the result of distortion in the sperm library.

**Figure 3.**
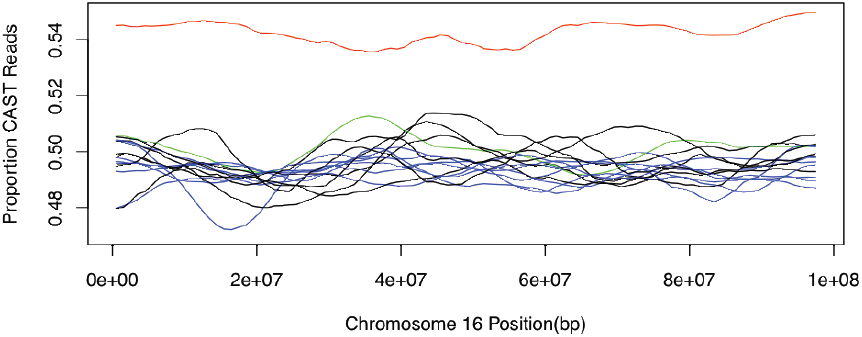
Proportion of informative reads that are derived from the CAST genome across chromosome 16. CW10’s liver sample is shown in red, and CW10’s sperm sample is shown in green. All other CW libraries are represented in black for liver and in blue for sperm.

Through simulation, we ensured that we have sufficient statistical power, given our experimental design and data quality, to detect segregation distortion if it is indeed occurring in hybrid males. We found that we have 50% power to detect SD to approximately 0.014, or 1.4% (this number reflects the positive or negative deviation from the null expectation, 0.5, at α = 0.001) if distortion affects CW and WC males equally (Figure 4). In other words, we have 50% power to detect distortion that is greater than 51.5% or less than 48.5%. If there is directionality to the distortion effect (*i.e.* only CW or only WC males experience SD), we have 50% power to detect distortion of 0.016 for CW males and 0.018 for WC males (at α = 0.001). This slight difference in power based on cross direction likely reflects differences in sequencing depth between WC and CW sperm and liver samples. It is also important to note that because read mapping and sequencing, as well as divergence between the CAST and WSB strains and their divergence from the reference genome, are non-uniform across the genome, different regions of the genome will differ slightly in power to detect distortion.

**Figure 4.**
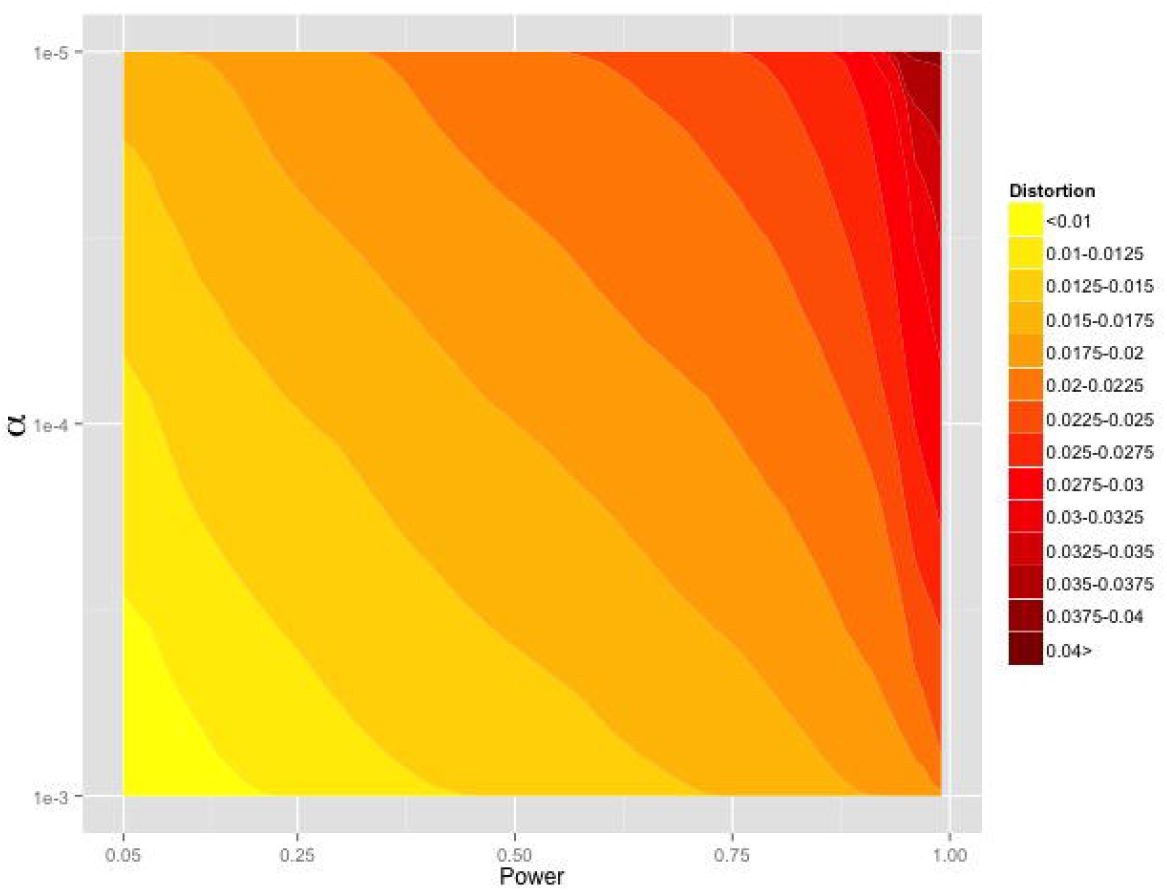
Minimum level of distortion that could be detected given a specified significance threshold (α, y-axis), and desired power (x-axis).

## Discussion

Elucidating the genetic mechanisms underlying species formation is a central goal of evolutionary biology. Although there has been progress in identifying the genetic basis of reproductive isolation in a few elegant instances (*e.g.* [36–38]), including several in *Drosophila* (*e.g.* [39,40]), it is unclear how general these results are. For example, SDs contribute to reproductive isolation in some young Drosophila species pairs ([13–15]) but here, to our surprise, we find no evidence for segregation distortion between two nascent species of mouse, *M. m. castaneus/M. m. domesticus,* despite strong experimental power.

Our conclusion must be qualified to some degree. SDs are generally classified as either gamete disablers or gamete killers depending on their mode of action (reviewed in [7,8]). We expect to detect gamete killers with our approach since their victims may not be present in the epididymides, or, if present, these sperm would not be captured in our stringent swim up assay. Our ability to detect gamete- disablers, however, depends on the specific mechanism by which these genetic elements act. If the motility or longevity of a sperm cell is sufficiently impaired, it is likely that this sperm would fail to swim into solution and remain motile over the course of the assay, but if the distortion effect has a very subtle effect on motility or impairs function later in the sperm life cycle (*e.g.* by causing a premature acrosome reaction), it is unlikely that our method could detect these effects. Thus, although gamete killers are not prevalent sources of DMIs in these subspecies, we cannot completely exclude the possibility that gamete disablers contribute to *M. musculus* species formation. However, it is worth nothing that disablers cannot explain the reported observation of increased apoptosis of sperm cells in hybrid males [26].

Conventional methods of detecting SDs (*i.e.* genotyping progeny) are usually statistically underpowered and thus unable to detect even modest distortion effects. Moreover, requiring the presence of offspring from F_1_ hybrids unavoidably conflates viability, gamete competition, and segregation distortion effects. By contrast, our simulations demonstrate that by sequencing high quality gametes from individual hybrid males and comparing allele ratios in these gametes to those of somatic tissues, we have excellent power to detect even relatively weak SDs, of less than two percent. In support of this point, we successfully detected an aneuploidy event that resulted in a four percent difference in allele frequencies relative to expectations within only a single biological replicate. Nonetheless, we found little evidence that SDs are active in F_1_ hybrid males, which indicates that segregation distortion (*i.e.* gamete killing) is not a primary contributor to reduced F_1_ male fertility in these subspecies.

Because our method of determining the allele ratios in bulk preparations of viable gametes relative to somatic tissues is very general, we expect that it will be useful in a wide variety of systems for an array of questions. Provided one can accurately phase the diploid genome of an individual, by *e.g.* using complete parental genotype data when inbred strains are not available, it is straightforward to apply this method to assay segregation distortion in a wide variety of taxa (including humans). Thus, we are now well positioned to survey the prevalence of segregation distortion both within and between a diversity of species. This approach also allows segregation distortion to be weighed against other possible sources of DMIs that may occur during spermatogenesis, oogenesis, fertilization, or embryogenesis, but that leaves an identical signature to SD in conventional cross-based experiments. Furthermore, because SDs can increase in frequency in populations despite deleterious consequences for the host, these selfish genetic elements may also be an important source of disease alleles. For example, it has been suggested that SDs contribute to the perpetuation of split-hand/split-foot disease [41], retinal dystrophy [42] and Machado-Joseph disease [43] in humans. Hence, the method introduced here has the potential to improve our understanding of disease evolution in addition to the contribution of SDs to the evolution of reproductive isolation between diverging lineages.

While segregation distorters may be an important mechanism of speciation in *Drosophila* and crop plants, efforts to detect SD in other diverging lineages—especially studies with high statistical power—have been limited. We find that at least in *M. m. castaneus/M. m. domesticus* hybrids, segregation distorters are not measurable contributors to observed infertility in F1 hybrid males, despite strong statistical power to detect them, suggesting that reduced hybrid male fertility in these nascent species is attributable to other underlying genetic causes. Further studies, using the novel approach developed here will provide a powerful way to gain more comprehensive understanding of the role of SDs within and between populations.

## Acknowledgements

The authors declare no conflicts of interest. We thank Tim Sackton, Julien Ayroles, Brian Arnold, Shelbi Russell, Brant Peterson, Marjorie Lienard, and Heidi Fisher for helpful comments and suggestions, and Catherine Dulac and Stacey Sullivan for assistance with mouse crosses. RCD was supported by a Harvard Prize Graduate Fellowship and currently a UCB Chancellor’s Postdoctoral Fellowship; EJP was supported by an NSF Predoctoral Fellowship.

## Author Contributions

RCD, EJP, DH, and HEH conceived and designed experiments. RCD, EJP, and HEH wrote the manuscript. RCD and EJP performed experiments. RCD analyzed the data.

---

Supplemental Text S1. Supplemental methods describing quality control steps to ensure samples are not contaminated or mislabeled.

Supplemental Table S1. List of genomic windows excluded from all downstream analyses due to detection of individual libraries with unusually high depth.

Supplemental Table S2. Quality control results for the quantity of reads in each library derived from the *Y* chromosome, *X* chromosome, and mtDNA.

Supplemental Table S3. Alignment simulation results showing the relationship between the reported mapping quality for a read and its probability of correct assignment to the genomic location from which it was derived.

